# BDNF induces its own release to mediate presynaptic plasticity

**DOI:** 10.1101/2021.12.30.474558

**Authors:** Coralie Berthoux, Kaoutsar Nasrallah, Teresa A. Milner, Pablo E. Castillo

**Affiliations:** Dominick P. Purpura Department of Neuroscience, Albert Einstein College of Medicine, Bronx, NY10461, United States; Feil Family Brain and Mind Research Institute, Weill Cornell Medicine, New York, NY10065, United States; Department of Psychiatry and Behavioral Sciences, Albert Einstein College of Medicine, Bronx, NY10461, United States

## Abstract

The brain-derived neurotrophic factor (BDNF) and its effector Tropomyosin receptor kinase B (TrkB) mediate diverse forms of activity-dependent synaptic plasticity implicated in learning, neural circuit refinement, and brain diseases, including epilepsy and mood disorders. Here, we report that activity-dependent release of presynaptic BDNF elicits the release of postsynaptic BDNF in a TrkB- and calcium-dependent manner. This BDNF-induced BDNF release was required for the induction of presynaptic long-term potentiation (LTP) of excitatory transmission in the mouse dentate gyrus. Tonic and phasic activity of presynaptic type-1 cannabinoid receptors suppressed BDNF release and dampened LTP, while exposure to enriched environment elicited BDNF-mediated LTP. In addition to mediating presynaptic plasticity, BDNF-induced BDNF release could be an important mechanism in synaptic stabilization during the maturation and refinement of neuronal connections.

**One-Sentence Summary:** The brain-derived neurotrophic factor induces its own release to mediate long-lasting increase in neurotransmitter release.

---

The mechanisms underlying experience-dependent synaptic plasticity, such as long-term potentiation (LTP), are remarkably diverse and not fully understood (*1, 2*). The brain-derived neurotrophic factor (BDNF) has been proposed as a key regulator of synaptic efficacy, neural connectivity, and experience-dependent plasticity (*3-6*). Consistent with these roles, BDNF is implicated in neural development, learning and memory (*3, 7-9*), as well as brain disorders (*10*). In the adult brain, BDNF mainly signals through the Tropomyosin receptor kinase B (TrkB) which can be located at presynaptic and postsynaptic sites (*3, 9*). BDNF is a soluble peptide stored in dense core vesicles and is released upon activity in a calcium- and SNARE-dependent manner (*11, 12*). However, it remains unclear how exactly BDNF/TrkB signaling mediates synaptic plasticity, including the kind of activity required to release BDNF and induce experience-dependent synaptic plasticity, the source of BDNF, and whether and how BDNF release is regulated (*4, 13*).

The dentate gyrus (DG), a key entry area of the hippocampus, expresses high levels of BDNF (*11, 14, 15*). Within the DG, two excitatory neurons, dentate granule cells (GCs) and hilar mossy cells (MCs) are reciprocally connected, thereby establishing a recurrent excitatory circuit (*16*). GCs send their axons to MCs, and MCs send their axons to the inner molecular layer (IML) to excite the proximal dendrites of GCs (*17, 18*). This associative GC→MC→GC circuit is believed to play key roles in DG-dependent learning (*19-21*). Dysregulation of this circuit is implicated in epilepsy (*18*). Remarkably, BDNF expression is particularly high in the IML (*11, 14, 15*), and TrkB is strongly expressed in both GC layer (GCL) and IML (*22-24*). Repetitive activation of MC axons induces robust LTP of MC-GC synaptic transmission (MC-GC LTP) that requires postsynaptic TrkB activation and is presynaptically expressed as a long-lasting increase in glutamate release (*25*). Unlike most forms of LTP at glutamatergic synapses, induction of MC-GC LTP does not require NMDA receptor activation or activation of any other ionotropic or metabotropic glutamate receptor (*25, 26*), suggesting that the release of BDNF, rather than glutamate, triggers MC-GC LTP. However, the source of BDNF and the release mechanism during activity-dependent MC-GC LTP remain undetermined. Intriguingly, MC axon terminals express high levels of type-1 cannabinoid receptors (CB_1_Rs) (*27-29*), whose activation by endocannabinoids (eCB) suppresses glutamate release (*30*). Whether and how eCB signaling controls BDNF release is unknown.

## Presynaptic BDNF release triggers LTP and is suppressed by cannabinoid signaling

MC-GC LTP requires postsynaptic BDNF/TrkB signaling (*25*) but the source of BDNF that mediates this plasticity is unclear. To test whether BDNF released from MC axons is required for induction, we selectively and conditionally removed *Bdnf* from MCs. Taking advantage of the MC commissural projections, we injected a mix of Cre recombinase-containing AAV (AAV5.CamKII.Cre-mCherry) and Cre-dependent ChiEF into one DG of *Bdnf*^fl/fl^ (Pre *Bdnf* cKO) or wild-type (WT, control) mice, and we analyzed the contralateral DG. This approach allowed us to optically activate Cre-positive (i.e., Pre *Bdnf* cKO in *Bdnf*^fl/fl^ mice) MC axon terminals in the IML (Fig. 1A–B). Paired-pulse ratio (PPR) and coefficient of variation (CV) were unchanged, suggesting that presynaptic *Bdnf* deletion did not significantly affect basal MC-GC presynaptic function (Fig. 1C). However, MC-GC LTP was abolished in Pre *Bdnf* cKO mice as compared to controls (Fig. 1D). In contrast, bath application of the adenylyl cyclase activator forskolin (50 μM for 10 min) potentiated MC-GC synaptic transmission in Pre *Bdnf* cKO mice similar to controls (Fig. 1E), consistent with the notion that presynaptic PKA activation downstream BDNF/TrkB signaling is necessary and sufficient for MC-GC LTP (*25*). Together, these findings indicate that the lack of MC-GC LTP was likely due to the absence of presynaptic BDNF in MC terminals.

**Figure 1.**
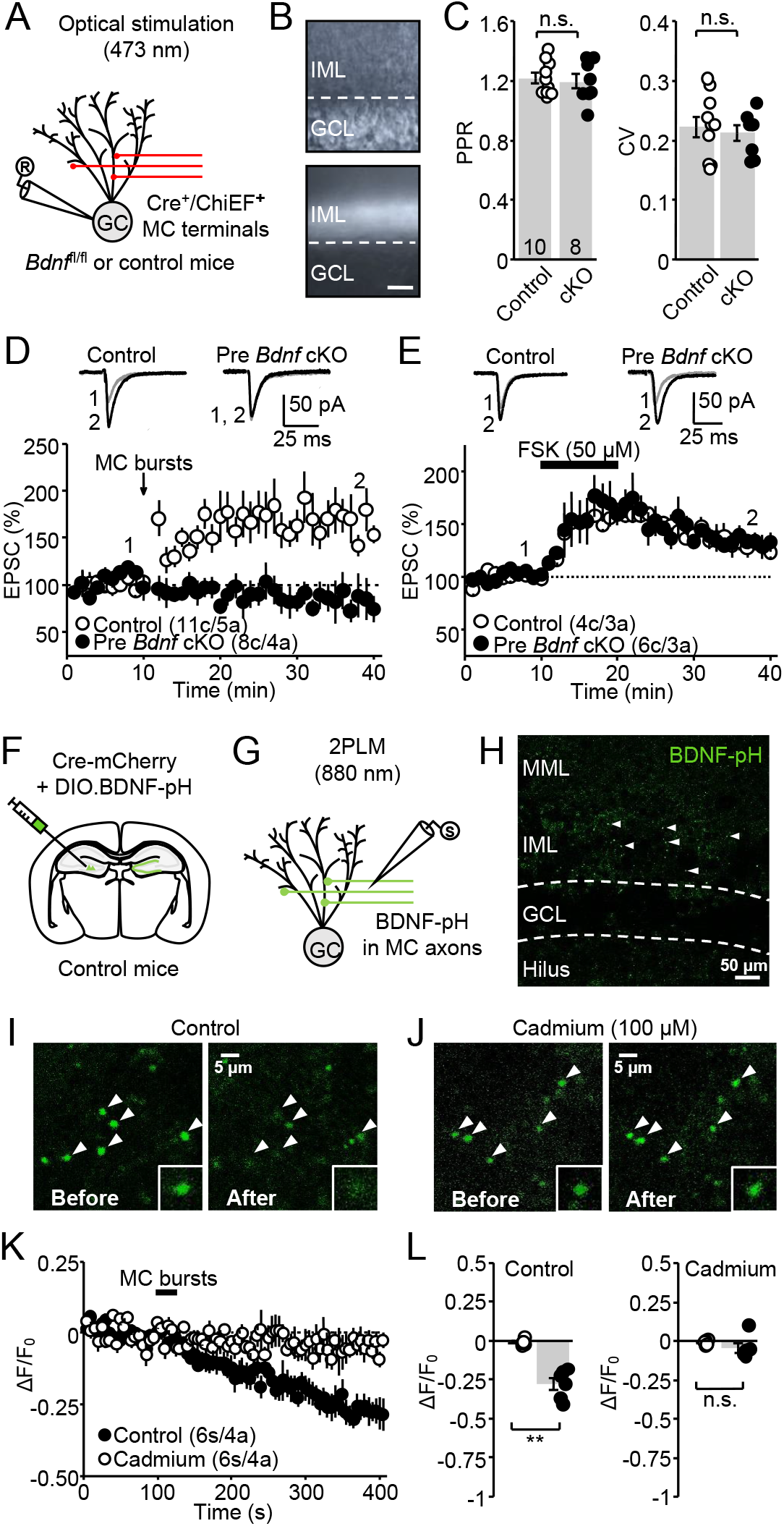
Presynaptic BDNF expression and release is required for MC-GC LTP. **(A)** Schematic diagram illustrating the optical stimulation of ChIEF-expressing MC axons and the recording configuration of GCs in contralateral hippocampal slices. A mix of AAV5.CaMKII.Cre-mCherry and AAV-DJ.FLEX.ChIEF-tdTomato was injected in the hilus of 3–4 w.o. *Bdnf*^fl/fl^ (Pre *Bdnf* cKO) or WT (control) mice. Whole-cell patch-clamp recordings were performed from GCs in contralateral DG. **(B)** Infrared/differential interference contrast (*top*) and fluorescence (*bottom*) images showing ChIEF-tdTomato was selectively expressed in the IML of contralateral DG of control and Pre *Bdnf* cKO mice. **(C)** Summary plots showing that basal PPR and CV were unchanged in Pre *Bdnf* cKO mice as compared to controls (PPR Control: 1.2 ± 0.04, n = 10; PPR cKO: 1.2 ± 0.05, n = 8; Control vs cKO, p = 0.7217, unpaired t-test; CV Control: 0.23 ± 0.02, n = 10; CV cKO: 0.21 ± 0.01, n = 8; Control vs cKO, p = 0.6343, unpaired t-test). **(D–E)** Optically-induced LTP was abolished in Pre *Bdnf* cKO mice (Control: 164 ± 10.4% of baseline, n = 11, p < 0.001, paired t-test; Pre *Bdnf* cKO: 86 ± 11.8% of baseline, n = 8, p = 0.1519, paired t-test; Control vs cKO, p < 0.001, unpaired t-test), but not FSK-induced LTP (Control: 133 ± 8.1% of baseline, n = 4, p < 0.05, paired t-test; Pre *Bdnf* cKO: 143 ± 6.4% of baseline, n = 6, p < 0.01, paired t-test; Control vs cKO, p = 0.3441, unpaired t-test). Vertical arrows correspond to the burst-stimulation of MC axons (5 pulses, 100 Hz, repeated 50 times every 0.5 s; light pulse duration 0.5–2 ms). Representative traces (1) and (2) correspond to the areas (1) and (2) in the summary time-course plots. For electrophysiology, data are represented as mean ± SEM. Number of cells and mice are showed between parentheses. n.s., not significant. **(F)** Schematic diagram illustrating the injection of a mix of AAV5.CaMKII.Cre-mCherry and AAV-DJ.DIO.BDNF-pHluorin (BDNF-pH) in the hilus of 3–4 w.o. WT (control) mice. **(G)** Experimental design of combined electrophysiology and 2-photon live imaging (2PLM) in acute hippocampal slices. Excitation wavelength employed was 880 nm. Extracellular field excitatory post-synaptic potentials (fEPSPs) were monitored using a stimulating pipette placed in the IML. **(H)** 2-photon image showing BDNF-pH was selectively expressed in the IML of contralateral DG (i.e., commissural MC axon terminals). Here and in all figures, arrowheads indicate BDNF-pH puncta. **(I–J)** Images showing changes in BDNF-pH fluorescence intensity after burst electrical stimulation of MC axon terminals in normal ACSF (**I**) or in the presence of VGCC inhibitor cadmium (100 μM, **J**). Here and in all figures, bottom right insets show a representative bouton at 1.8x magnification. **(K)** Time-course summary plot showing the average fractional fluorescence changes (ΔF/F_0_) over time. **(L)** Quantification of the averaged responses before and 300–400 s following MC burst stimulation (Control: -0.28 ± 0.03, n = 6, p < 0.01, paired t-test; Cadmium: -0.04 ± 0.03, n = 6, p = 0.3749, paired t-test; Control vs Cadmium, p < 0.001, unpaired t-test). For imaging, data are represented as mean ± SEM. Here and in all figures, number of slices (s) and animals (a) are showed between parentheses. ** p < 0.01; n.s., not significant.

To demonstrate that BDNF is released from MC axons upon activity, we utilized a BDNF reporter (BDNF-pHluorin or BDNF-pH) (*31-33*). To restrict the expression of BDNF-pH in MC axon terminals, a mix of Cre recombinase-containing AAV (AAV5.CamKII.Cre-mCherry) and Cre-dependent BDNF-pH was injected into one DG of WT mice (Fig. 1F). Four weeks post-injection, we observed a high-level BDNF-pH expression in commissural MC axons of the contralateral IML (Fig. 1G–H). To assess activity-dependent BDNF release, we monitored changes in the fluorescence intensity of BDNF-pH puncta (1–3 μm), which represent BDNF-containing presynaptic boutons, using time-lapsed 2-photon microscopy of acute hippocampal slices (Fig. 1H). We found that repetitive activation of MC axons (MC bursts) that induces MC-GC LTP (5 pulses, 100 Hz, repeated 50 times every 0.5 s) significantly reduced BDNF-pH fluorescence intensity (Fig. 1I and 1K–L). BDNF requires calcium influx via VGCCs (*13, 31*). As expected, bath application of the non-selective VGCC blocker cadmium (100 μM) abolished the reduction in BDNF-pH fluorescence (Fig. 1J–L). These results indicate that repetitive activation of MC axons triggers BDNF release from axon terminals in a presynaptic calcium influx-dependent manner.

MC axon boutons express uniquely high levels of CB_1_Rs (*27-29*), which inhibit glutamate release and MC-GC LTP induction in a tonic (i.e., constitutively active, ligand-independent) and phasic (i.e., ligand-dependent) manner (*26*). Given that LTP induction requires BDNF/TrkB signaling (*25*) but not glutamate transmission (*26*), we tested whether presynaptic CB_1_Rs dampen LTP by suppressing BDNF release from MC axonal boutons. To this end, we pre-incubated acute hippocampal slices with the CB_1_R agonist WIN 55,212-2 (5 μM for 25 min), which was also perfused in the recording chamber. BDNF release following the MC-GC LTP induction protocol was suppressed by WIN (Fig. 2A and 2C–D), and this effect was prevented by the CB_1_R inverse agonist AM251 (5 μM) (Fig. 2B–D). These results indicate that activation of CB_1_R inhibits BDNF release from MC axons. In addition to this ligand-dependent effect via presynaptic CB_1_R activation, presynaptic BDNF release could also be under the control of tonic CB_1_R activity. To test this possibility, MC axons were stimulated with bursts known to induce a submaximal LTP (e.g., 30 Hz instead of 100 Hz bursts used in Fig. 1F–L) which is facilitated by blocking CB_1_R constitutive activity with AM251 (*26*). We found that blocking tonic CB_1_R activity with AM251 (5 μM) facilitated presynaptic BDNF release, as indicated by a large reduction in BDNF-pH fluorescence intensity as compared to controls (Fig. 2F–H). Low sensitivity of BDNF-pHluorin could account for the lack of change in fluorescence following submaximal LTP in absence of AM251 (Fig. 2E and 2G–H). Altogether, these results indicate that CB_1_R activity inhibits presynaptic BDNF release, thereby dampening MC-GC LTP induction.

**Figure 2.**
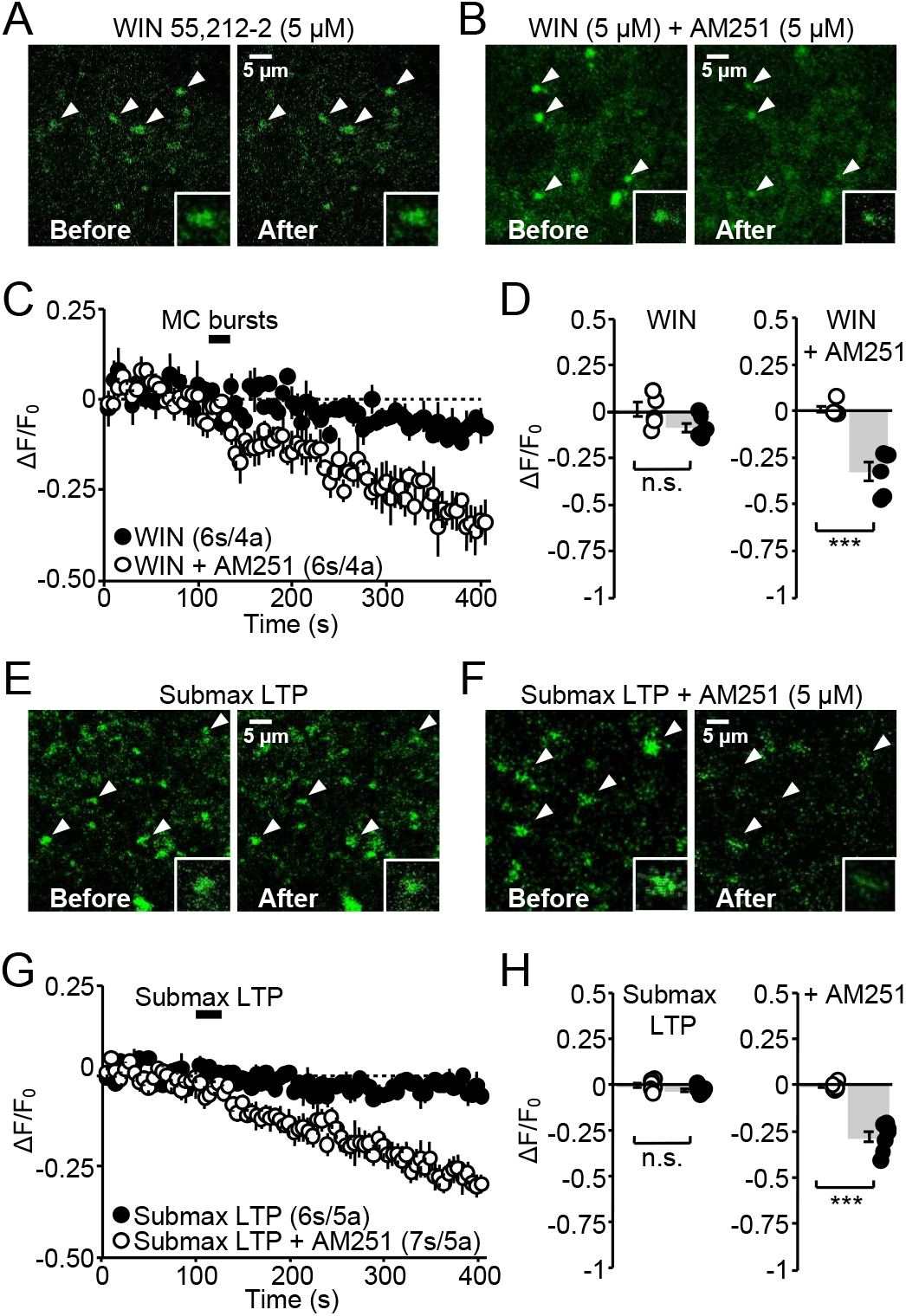
Presynaptic BDNF release is under the control of CB_1_R activity. **(A–B)** CB_1_R activation by WIN 55,212-2 (5 μM, pre-incubated for 25 min and perfused in the chamber) prevented changes of BDNF-pH fluorescence intensity following burst electrical stimulation of MC axon terminals. This effect was abolished in slices co-incubated with CB_1_R inverse agonist AM251 (5 μM, for 25 min). **(C)** Time-course summary plot showing the average fractional fluorescence changes (ΔF/F_0_) over time. **(D)** Quantification of the averaged responses 300–400 s following MC burst stimulation (WIN: -0.08 ± 0.02, n = 6, p = 0.0707, paired t-test; WIN+AM251: -0.32 ± 0.05, n = 6, p < 0.001, paired t-test; WIN vs WIN+AM251, p < 0.01, unpaired t-test). **(E–F)** A 30 Hz induction protocol was applied to induce submaximal LTP, and this protocol alone was not sufficient to induce significant BDNF release from MC axons. Inhibition of tonic CB_1_Rs by bath application of AM251 (5 μM, for 25 min) decreased BDNF-pH fluorescence intensity following submaximal LTP. **(G)** Time-course summary plot showing the average fractional fluorescence changes (ΔF/F_0_) over time. **(H)** Quantification of the averaged responses following 300–400 s MC burst stimulation (Submaximal LTP: -0.03 ± 0.01, n = 6, p = 0.1216, paired t-test; Submaximal LTP + AM251: -0.28 ± 0.03, n = 7, p < 0.01, paired t-test; Submaximal LTP vs Submaximal LTP + AM251, p < 0.001, unpaired t-test). Data are represented as mean ± SEM. *** p < 0.001; n.s., not significant.

### Postsynaptic TrkB activation promotes postsynaptic BDNF release and presynaptic LTP

BDNF is found in GC dendrites (*14*) from where it could be released to activate postsynaptic TrkB in an autocrine manner (*34*), or target presynaptic TrkB in a retrograde manner (*35, 36*). To test whether postsynaptic BDNF could also be involved in MC-GC LTP, we conditionally and selectively knocked out *Bdnf* from GCs by injecting AAV5.CamKII.Cre-mCherry (Post *Bdnf* cKO) or AAV5.CamKII.mCherry (control) in the upper GCL of *Bdnf*^fl/fl^ mice (Fig. 3A and fig. S1A), and we recorded MC-elicited EPSCs from mCherry-positive GCs. Deleting *Bdnf* from GCs had no significant effect on basal MC-GC PPR and CV (fig. S1B). Surprisingly, MC-GC LTP was abolished in Post *Bdnf* cKO (Fig. 3B), while forskolin-induced LTP was unaffected (Fig. 3C), consistent with cAMP/PKA acting downstream of BDNF/TrkB signaling (*25*). To test whether BDNF was present in dendrites or dendritic spines, we analyzed a previously characterized mouse line in which *Bdnf* coding sequence is fused to a C-terminal haemagglutinin (HA) epitope tag (*Bdnf*-HA) (*34*). Using electron microscopy and highly sensitive antibodies against HA-tag, we found HA-BDNF-immunoreactivity in the IML is localized to discrete patches and is present in axon terminals, which putatively arise from MC axons, as well as in dendrites and dendritic spines of GCs (fig. S2). To test whether burst stimulation of MC axons releases BDNF from GCs during the induction of MC-GC LTP, we measured postsynaptic BDNF release selectively by expressing BDNF-pH in the upper GCL of WT mice (Fig. 3D). Ten days post-injection we observed a high-level of BDNF-pH in the soma and proximal dendrites of GCs with Cre expression restricted to the upper GCL (fig. S3). MC burst stimulation significantly reduced BDNF-pH fluorescence intensity from GC proximal dendrites (Fig. 3E–G). Although BDNF release could result from postsynaptic firing (*13*), MC bursts are not sufficient to induce GC firing under our experimental conditions (i.e., in absence of GABA receptor antagonists) (*25*). Lastly, postsynaptic BDNF release was abolished by the TrkB antagonist ANA-12 (15 μM for 25 min) (Fig. 3E–G), revealing that postsynaptic BDNF release requires TrkB activation. Altogether, our findings indicate that presynaptic burst activity triggers postsynaptic BDNF release in a TrkB-dependent manner, and that release of postsynaptic BDNF is required for the induction of LTP at MC-GC synapses.

**Figure 3.**
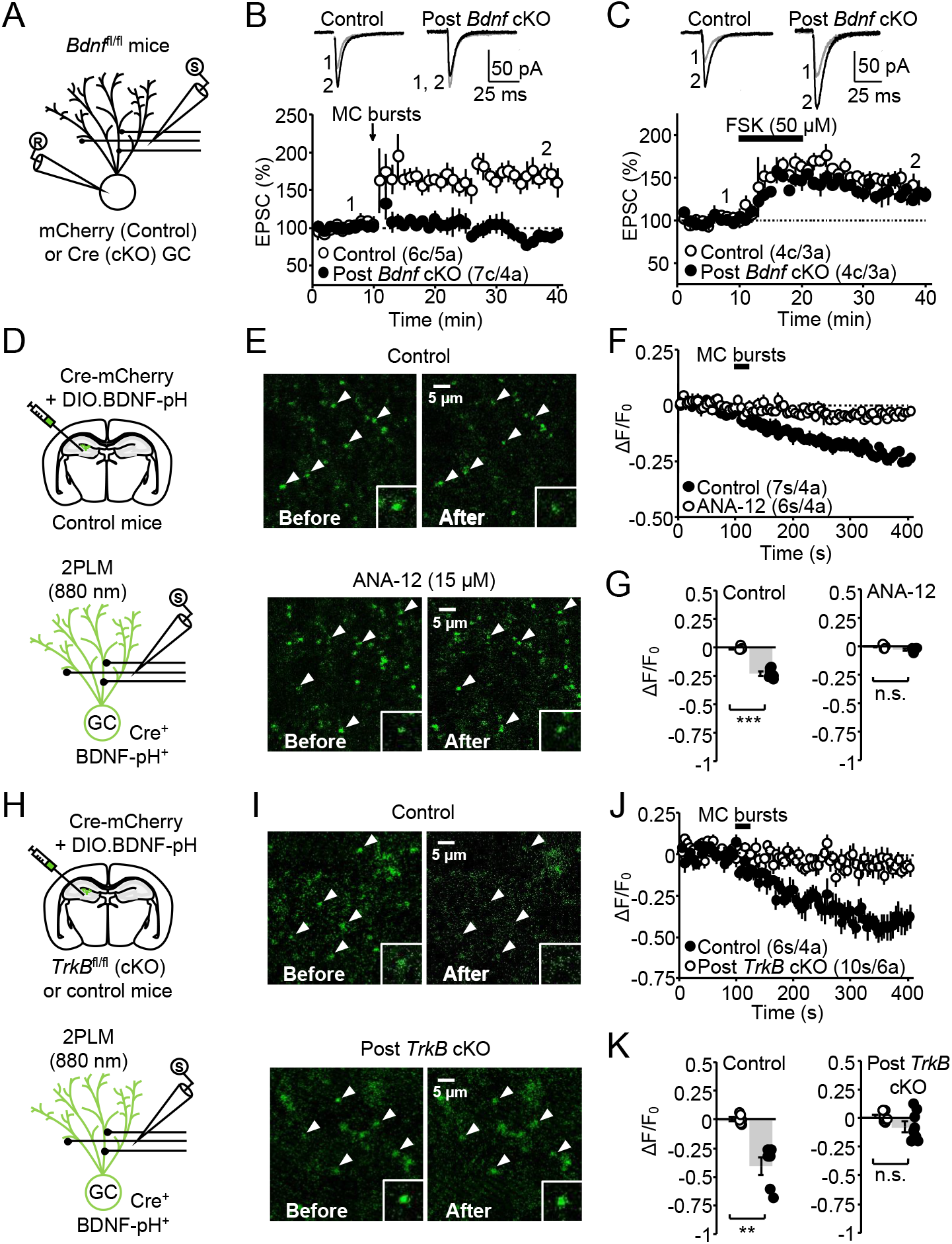
Postsynaptic BDNF is required for presynaptic LTP and its release is TrkB-dependent. **(A)**Schematic diagram illustrating the recording configuration of GCs from mCherry-positive GCs and the electrical stimulation of MC axons in ipsilateral hippocampal slices. AAV5.CaMKII.mCherry (control) or AAV5.CaMKII.Cre-mCherry (Post *Bdnf* cKO) was injected in the upper GCL of 3-4 w.o. *Bdnf*^fl/fl^ mice. Whole-cell patch-clamp recordings were performed from mCherry-expressing GCs. **(B–C)** Synaptically-induced LTP was abolished in Post *Bdnf* cKO mice (Control: 168 ± 13% of baseline, n = 6, p < 0.05, paired t-test; Post *Bdnf* cKO: 103 ± 9% of baseline, n = 7, p = 0.8682, Wilcoxon signed rank test; Control vs cKO, p < 0.01, Mann-Whitney’s U test), but not FSK-induced LTP (**C**) (Control: 142 ± 7% of baseline, n = 4, p < 0.01, paired t-test; Post *Bdnf* cKO: 138 ± 6% of baseline, n = 4, p < 0.01, paired t-test; Control vs cKO, p = 0.6959, unpaired t-test). Vertical arrow corresponds to the burst-stimulation of MC axons (5 pulses, 100 Hz, repeated 50 times every 0.5 s). Representative traces (1) and (2) correspond to the time points (1) and (2) in the summary time course plots. Data are presented as mean ± SEM. n.s., not significant. **(D)** Schematic diagram (*top*) illustrating the injection of a mix of AAV5.CaMKII.Cre-mCherry and AAV-DJ.DIO.BDNF-pHluorin (BDNF-pH) in the upper GC layer of 3–4 w.o. WT (control) mice. Experimental design of combined electrophysiology and 2PLM in acute hippocampal slices (*bottom*). Excitation wavelength employed was 880 nm. fEPSPs were monitored using a stimulating pipette in the IML. **(E)** Representative images showing changes in BDNF-pH fluorescence intensity in GC proximal dendrites, after burst electrical stimulation of MC axon terminals in normal ACSF (*top*) or in the presence of a TrkB inhibitor ANA-12 (*bottom*). (**F**) Time-course summary plot showing the average fractional fluorescence changes (ΔF/F_0_) over time. (**G**) Quantification of the averaged responses 300–400 s following MC burst stimulation (Control: -0.23 ± 0.01, n = 7, p < 0.001, paired t-test; ANA-12: -0.02 ± 0.01, n = 6, p = 0.0649, paired t-test; Control vs ANA-12, p < 0.001, unpaired t-test). **(H)** Schematic diagram illustrating the injection as in (D) in the upper GC layer of 3–4 w.o. *TrkB*^fl/fl^ (Post *TrkB* cKO) or WT (control) littermates. **(I)** Representative images as in (E) from Post *TrkB* cKO and control littermates, after burst electrical stimulation of MC axon terminals. **(J)** Time-course summary plot showing the average fractional fluorescence changes (ΔF/F_0_) over time. **(K)** Quantification of the averaged responses 300–400 s following MC burst stimulation (Control: -0.40 ± 0.02, n = 6, p < 0.01, Wilcoxon signed rank test; Post *TrkB* cKO: -0.07 ± 0.05, n = 10, p = 0.1404, paired t-test; Control vs cKO, p < 0.01, Mann-Whitney’s U test). Data are represented as mean ± SEM. *** p < 0.001; ** p < 0.01; n.s., not significant.

Given that activation of postsynaptic TrkB is necessary to induce MC-GC LTP (*25*), we directly measured BDNF release from *TrkB*-lacking GCs (Post *TrkB* cKO) by injecting AAV5.CamKII.Cre-mCherry alongside with Cre-dependent BDNF-pH into the DG of *TrkB*^fl/fl^ mice or WT (control) littermates (Fig. 3H). We found that burst stimulation of MC axons significantly reduced BDNF-pH fluorescence intensity in controls, but not in Post *TrkB* cKO mice (Fig. 3I–K), strongly suggesting that presynaptically released BDNF, by activating postsynaptic TrkB, promotes BDNF release from GC dendrites. BDNF could also activate presynaptic TrkB (*35-38*), thereby facilitating further BDNF release from MC axons. To test this possibility, we selectively and conditionally removed *TrkB* from MC axon terminals by injecting a mix of AAV5.CamKII.Cre-mCherry and Cre-dependent ChiEF into the DG of *TrkB*^fl/fl^ (Pre *TrkB* cKO) or WT (control) littermates (fig. S4A). This approach allowed us to optically activate Cre-positive (i.e., Pre *TrkB* cKO in *TrkB*^fl/fl^ mice) MC axon terminals in the IML of the contralateral hippocampus (fig. S4B–D). MC-GC LTP was normally induced in Pre *TrkB* cKO mice as compared to controls (fig. S4E), suggesting that presynaptic TrkB expressed on MC axons are not required for MC-GC LTP induction.

### BDNF is sufficient to induce BDNF release from GC proximal dendrites

To test whether BDNF is sufficient to induce postsynaptic BDNF release, we monitored BDNF-pH fluorescence intensity and bath applied a human recombinant BDNF (8 nM for 10 min). We found that exogenous BDNF application induced BDNF release from GC dendrites (Fig. 4A and 4D–E), and this effect was abolished in the presence of the TrkB antagonist ANA-12 (15 μM) (Fig. 4B and 4D–E). BDNF release from GC could result from postsynaptic firing (*13*). However, bath application of BDNF for 10 min did not trigger any GC firing under our recording conditions (no drugs in the bath) (fig. S5A and S5C), whereas subsequent blockade of inhibitory transmission by bath applying 100 μM picrotoxin robustly increased GC burst firing (fig. S5A–C) as expected (*25*). These results demonstrate that exogenous BDNF application triggers BDNF release from GCs in a TrkB-dependent but action potential–independent manner. Calcium release from internal stores is implicated in BDNF release (*11, 31, 32*), and TrkB can mobilize calcium internal stores via phospholipase C (PLC) (*39*). In addition, BDNF release can occur following calcium mobilization through activation of the sarcoplasmic reticulum Ca^2+^ (SERCA) ATPases (*40-42*). We measured BDNF release in the presence of the SERCA ATPase inhibitor CPA (30 μM) and found that BDNF-induced BDNF release was abolished (Fig. 4C and 4D–E). In addition, both the PLC inhibitor U73122 (5 μM) and CPA (30 μM) prevented MC-GC LTP (Fig. 4F), suggesting that TrkB activation engages PLC signaling and mobilizes calcium from internal stores to release BDNF. None of these inhibitors had a significant effect on basal synaptic transmission (Fig. 4G). Together, these findings indicate that BDNF released from MCs is necessary and sufficient to induce BDNF release from GCs via a TrkB-dependent calcium mobilization from internal stores.

**Figure 4.**
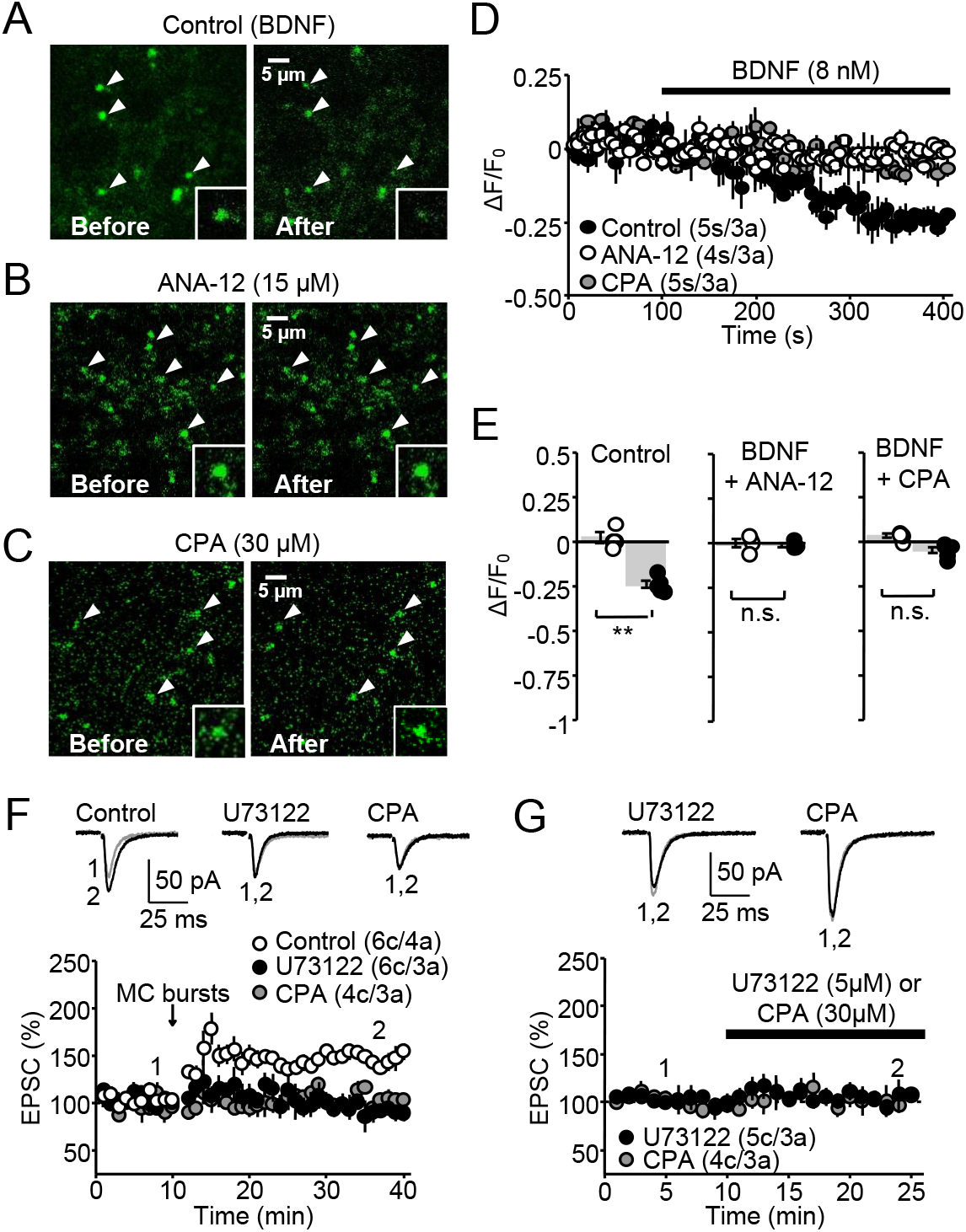
BDNF is sufficient to induce BDNF release by mobilizing calcium from internal stores. **(A–C)** 2-photon images showing changes in BDNF-pH fluorescence intensity in GC proximal dendrites, following bath application of BDNF (8 nM) in normal ACSF (control, **A**) or in the presence of a TrkB inhibitor ANA-12 (**B**) or the SERCA pump blocker cyclopiazonic acid CPA (**C**). **(D)** Time-course summary plot showing the average fractional fluorescence changes (ΔF/F_0_) over time. **(E)** Quantification of the averaged responses 300–400 s following BDNF bath application. BDNF (8 nM) decreased BDNF-pH fluorescent intensity (Control: -0.24 ± 0.02, n = 5, p < 0.01, paired t-test). This effect was abolished in the presence of 15 μM ANA-12 (BDNF + ANA-12: -0.01 ± 0.01, n = 4, p = 0.6538, paired t-test) or 30 μM CPA (BDNF + CPA: -0.05 ± 0.02, n = 5, p = 0.0691, Wilcoxon signed rank test). **(F)** MC-GC LTP was abolished in the presence of PLC inhibitor U73122 (5 μM, pre-incubated 1 h) or the inhibitor of the SERCA pump blocker cyclopiazonic acid CPA (30 μM) as compared to naïve slices (Control: 149 ± 5% of baseline, n = 6, p < 0.001, paired t-test; U73122: 103 ± 8% of baseline, n = 6, p = 0.9687, paired t-test; CPA: 104 ± 4% of baseline, n = 4, p = 0.4390, paired t-test; Control vs U73122: p < 0.001, Control vs CPA: p < 0.001, U73122 vs CPA: p = 0.9994, one-way ANOVA followed by a Tukey’s *post hoc* test). **(G)** Bath application of U73122 (5 μM) or CPA (30 μM) did not affect basal synaptic transmission (U73122: 103 ± 6% of baseline, n = 5, p = 0.8414, paired t-test; CPA: 104 ± 9% of baseline, n = 4, p = 0.7337, paired t-test). Data are represented as mean ± SEM. Number of slices (s), cells (c) and animals (a) are showed between parentheses. ** p < 0.01; n.s., not significant.

### Exposure to enriched environment strengthens MC-GC synaptic transmission in a BDNF-dependent manner

BDNF is commonly implicated in the neuronal and synaptic changes associated with exposure to an enriched environment (EE) (*43*). There is good evidence that EE and voluntary exercise improve hippocampus-dependent learning (*43-45*), which likely relies on diverse mechanisms, including activity-dependent synaptic plasticity and increase in BDNF/TrkB signaling (*46-48*). Given the essential role of BDNF/TrkB in MC-GC plasticity, we examined whether EE exposure compared to a standard cage (home cage, HC) could potentiate MC-GC synaptic transmission in a BDNF-dependent manner. We found that in EE mice MC-GC synaptic transmission assessed by minimal stimulation in the IML (Fig. 5A–B) was characterized with a significant increase in success rate, an increase in efficacy (i.e., mean EPSC amplitude including failures), but no change in potency (i.e., mean EPSC amplitude excluding failures) (Fig. 5C). In addition, PPR and CV were decreased in EE mice compared to HC mice (Fig. 5D), indicative of an increase in evoked neurotransmitter release. Remarkably, MC-GC LTP was occluded in *ex-vivo*, acute hippocampal slices obtained from EE as compared to HC mice (Fig. 5E). In addition, we found that 1-hour exposure to EE was sufficient to reduce PPR and CV (fig. S6A–C) and occlude MC-GC LTP (fig. S6D). Together, these results strongly suggest that EE exposure induced presynaptic strengthening of MC-GC transmission likely due to *in vivo* induction of MC-GC LTP.

**Figure 5.**
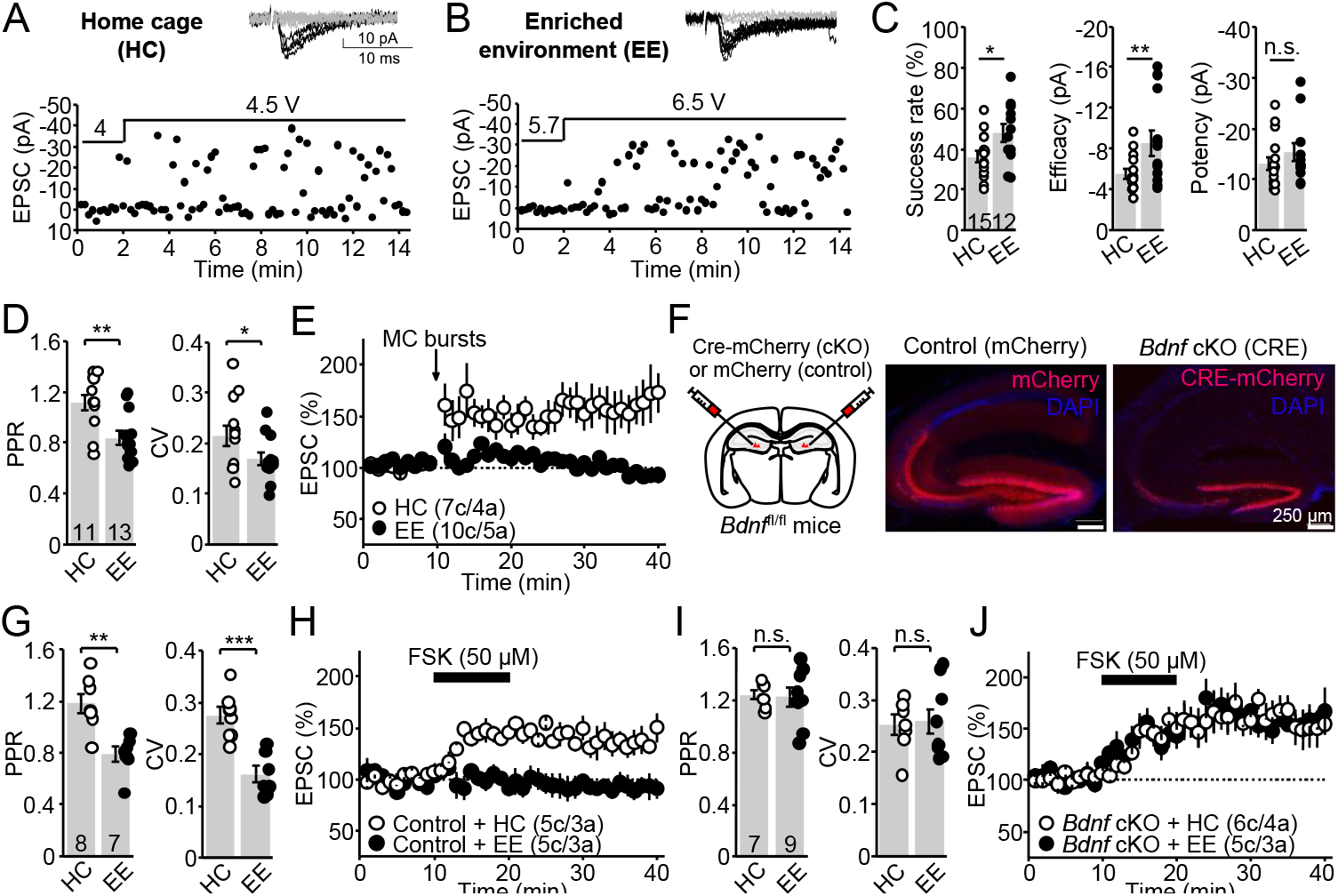
Exposure to enriched environment strengthens MC-GC synapses in a BDNF-dependent manner. **(A-B)** Representative experiments showing synaptic responses evoked by minimal stimulation in the IML of WT mice exposed to HC (**A**) or EE (**B**). Sample traces (*top*) and time-course plot (*bottom*). **(C)** Summary plots showing that EE mice are characterized by a significant increase in success rate (HC: 36 ± 2.8%, n = 15; EE: 48 ± 4%, n = 12; p < 0.05, unpaired t test), an increase in efficacy (HC: -5.5 ± 0.5 pA, n = 15; EE: -8.5 ± 1.2 pA, n = 12; p < 0.01; Mann-Whitney’s U test), and no change in potency (HC: -13.2 ± 1.4 pA, n = 15; EE: -15.4 ± 1.7 pA, n = 12, p = 0.17; Mann-Whitney’s U test). **(D)** Summary plots showing that PPR and CV were decreased in mice exposed to EE as compared to control HC mice (PPR HC: 1.1 ± 0.07, n = 11; EE: 0.8 ± 0.05, n = 13; HC vs EE, p < 0.01, unpaired t-test; CV HC: 0.22 ± 0.02, n = 11; EE: 0.16 ± 0.01, n = 13; HC vs EE, p < 0.05, unpaired t-test). **(E)** Synaptically-induced LTP was occluded in mice exposed to EE for 2 weeks as compared to controls (HC: 159 ± 19% of baseline, n = 7, p < 0.05, paired t-test; EE: 93 ± 6% of baseline, n = 10, p = 0.0880, paired t-test; HC vs EE: p < 0.05, unpaired t-test). (**F**) Schematic diagram (*left*) illustrating the injection of AAV5.CaMKII.mCherry (control) or AAV5.CaMKII.Cre-mCherry (*Bdnf* cKO) in the DG of 3–4 w.o. *Bdnf*^fl/fl^ mice. Confocal images show mCherry and Cre-mCherry were expressed in both GCs and MCs. **(G)** Summary plots showing that PPR and CV were reduced in control mice exposed to EE as compared to HC (PPR HC: 1.2 ± 0.07, n = 8; EE: 0.8 ± 0.06, n = 7; HC vs EE, p < 0.01, unpaired t-test; CV HC: 0.28 ± 0.02, n = 8; EE: 0.16 ± 0.02, n = 7; HC vs EE, p < 0.001, unpaired t-test). **(H)** Forskolin (FSK)-induced LTP was abolished in EE-exposed mice as compared to controls (HC: 138 ± 8% of baseline, n = 5, p < 0.05, paired t-test; EE: 92 ± 6% of baseline, n = 5, p = 0.2406, paired t-test; HC vs EE, p < 0.01, unpaired t-test). **(I)** No changes in PPR and CV were observed in *Bdnf* cKO mice exposed to HC or EE (PPR HC: 1.2 ± 0.04, n = 7; EE: 1.2 ± 0.07, n = 9; HC vs EE, p = 0.8609, unpaired t-test; CV HC: 0.25 ± 0.03, n = 7; EE: 0.26 ± 0.03, n = 9; HC vs EE, p = 0.8441, unpaired t-test). **(J)** Forskolin (FSK)-induced LTP was intact in *Bdnf* cKO mice after HC or EE exposure (HC: 161 ± 15% of baseline, n = 6, p < 0.05, paired t-test; EE: 148 ± 10% of baseline, n = 5, p < 0.01, paired t-test; HC vs EE, p = 0.5196, unpaired t-test). Data are represented as mean ± SEM. *** p < 0.001; ** p < 0.01; * p < 0.05; n.s., not significant.

Lastly, we sought to determine whether EE-induced potentiation of MC-GC synaptic transmission also requires BDNF/TrkB signaling, as observed in acute hippocampal slices. We therefore injected AAV5.CamKII.Cre-mCherry (*Bdnf* cKO) or AAV5.CamKII.mCherry (control) in the DG of *Bdnf*^fl/fl^ mice. One week post-surgery recovery, mice were placed in EE or HC for two weeks, and MC-GC EPSCs were then recorded from mCherry-positive GCs in acute hippocampal slices. Post-hoc analysis confirmed that Cre-mCherry and mCherry were expressed in both MCs and GCs (Fig. 5F). Consistent with our previous results, control mice exposed to EE showed reduced PPR and CV (Fig. 5G), and a failure in forskolin-induced LTP (Fig. 5H). In contrast, PPR and CV remained unchanged in *Bdnf* cKO mice exposed to EE compared to HC (Fig. 5I), suggesting a failure in EE-mediated strengthening of MC-GC synaptic transmission. Because cAMP/PKA signaling is required for MC-GC LTP downstream BDNF/TrkB (*25*), we then examined whether cAMP/PKA activation could still induce MC-GC LTP in *Bdnf* cKO mice using forskolin. Forskolin (50 μM for 10 min) induced comparable MC-GC potentiation in *Bdnf* cKO mice exposed to EE and HC (Fig. 5J). Taken together, our results reveal that EE exposure likely induces BDNF/TrkB-dependent LTP at MC-GC synapses.

### BDNF-induced BDNF release gates presynaptic plasticity

In this study, we report that endogenous BDNF induces its own release to mediate synaptic plasticity in the mature brain, and that this positive feedback mechanism is under the control of cannabinoid signaling (fig. S7). While common triggers of BDNF release include postsynaptic depolarization and the activation of metabotropic and ionotropic glutamate receptors (e.g., mGluR1/5 and NMDA receptors, respectively) (*3, 13, 49*), MC-GC LTP can be induced in the absence of postsynaptic depolarization, glutamate release or activation of glutamate receptors (*25, 26*). BDNF-induced BDNF release likely plays an instructive role for the induction of presynaptic long-term plasticity, presumably by locally increasing BDNF above critical concentrations. Our findings are consistent with an early study reporting that exogenous application of neurotrophins induces BDNF release in both cultured neurons and acute hippocampal slices, and this release depends on activation of neurotrophin receptors from the Trk family –as indicated by the kinase inhibitor K252a– and mobilization of calcium from internal stores (*50*). However, we here provide direct evidence that endogenous release of BDNF from presynaptic terminals upon repetitive activity induces postsynaptic BDNF release. We cannot discard that postsynaptic BDNF release promotes further release of BDNF in an autocrine manner (*34*). Moreover, we show that presynaptic removal of TrkB did not affect MC-GC LTP, making unlikely that BDNF may act as a retrograde messenger in the induction of this presynaptically expressed form of plasticity. BDNF-induced BDNF release could also play important roles in the maturation and refinement of neuronal connections. For example, it could act as positive feedback required for synaptic stabilization (*50*), and axon initiation and growth during development (*51*).

BDNF expression and release are highly regulated processes (*3, 13*). Consistent with the strong expression of CB_1_Rs in MC axon terminals (*27-29*), we found that tonic and phasic CB_1_R activity negatively regulates presynaptic BDNF release from MC axons, revealing an undescribed role of endocannabinoid signaling in suppressing neuropeptide release. Like for glutamate release (*26*), the CB_1_R-mediated reduction of presynaptic BDNF release is likely due to a βγ subunit-mediated suppression of presynaptic calcium influx, although a potential contribution due to cAMP/PKA reduction via the α_i/o_ subunit cannot be discarded given that BDNF secretion is reportedly facilitated by intracellular cAMP (*51-53*). By reducing presynaptic BDNF release from MC axons, CB_1_Rs can control MC-GC synaptic strength in a long-term manner. This action may contribute to DG-dependent behaviors that rely on a sparse GC activity. Moreover, some of the cannabinoid-mediated effects on brain development (*54*) could be due to the suppression of BDNF release. Because BDNF promotes temporal lobe epilepsy (*55, 56*), and BDNF/TrkB-mediated MC-GC LTP facilitates early seizure activity (*57*), the CB_1_R-mediated reduction of presynaptic BDNF release could be a mechanism by which cannabinoids may dampen epileptic activity.

Enriched environment and exercise are known to promote BDNF expression in different brain areas, including the DG (*46-48*), and improve cognition (*43*). We found that exposure to enriched environment strengthens MC-GC synaptic transmission in a presynaptic manner. In addition, knocking out *Bdnf* from GCs abolished enriched environment-induced MC-GC synaptic strengthening, strongly suggesting that *in vivo* MC-GC LTP induction requires BDNF/TrkB signaling as described in acute hippocampal slices (this study and *25*). Molecular tools that specifically target MCs or GCs in a subcellular compartment manner will be required to further characterize the precise involvement of BDNF arising from MCs and GCs *in vivo*. Together, the robust CB_1_R-mediated reduction of presynaptic BDNF release and MC-GC synaptic strength may play important roles in DG-dependent computations, such as pattern separation and spatial memory. Given the crucial role of BDNF/TrkB signaling in epilepsy, depression, and anxiety (*9, 10*), BDNF-induced BDNF release could be a target for the treatment of these brain disorders.

## List of Supplementary Materials

Materials and Methods

Figs. S1 to S7

References (*58*)

## Acknowledgments

We thank Drs. P. Kaeser (Harvard Medical School) and H. Park (Korea Brain Research Institute) for providing pAAV-FLEX-ChIEF-tdTomato and pAAV-DIO-BDNF-pHluorin, respectively. We also thank Dr. Lisa Monteggia (University of Texas, Southwestern Medical Center) for providing floxed-*TrkB* and floxed-*Bdnf* mice. We thank the Neuroanatomy Electron Microscopy Core at Weill Cornell Medicine for assistance with the EM study. Confocal images were obtained at the Einstein Imaging Core (supported by The Rose F Kennedy Intellectual Disabilities Research Center U54 HD090260, and shared instrument grant 1S10OD25295 to Konstantin Dobrenis).

## Funding

National Institutes of Health R01-MH125772 (PEC)

National Institutes of Health R01-NS113600 (PEC)

National Institutes of Health R01-NS115543 (PEC)

National Institutes of Health R01-MH116673 (PEC)

National Institutes of Health R01-DA 008259 (TAM)

Junior Investigator Neuroscience Research Award -Einstein (CB)

American Epilepsy Society Postdoctoral Award (KN)

## Author Contributions

Conceptualization: CB, PEC

Methodology: CB, TAM

Investigation: CB, KN, TAM

Visualization: CB, PEC

Funding acquisition: CB, KN, TAM, PEC

Project administration: PEC

Supervision: PEC

Writing – original draft: CB, PEC

Writing – review & editing: CB, KN, TAM, PEC

## Data and materials availability

All data are available in the main text or the supplementary materials.

