## Supplemental Materials for "BDNF induces its own release to mediate presynaptic plasticity"

**This PDF file includes:**

Materials and Methods
Figs. S1 to S7
References (*58*)

### Materials and methods

#### Experimental model and subject details

C57BL/6J (Charles River), floxed-*Bdnf* (*Bdnf*$^{fl/fl}$) and floxed-*TrkB* (*TrkB*$^{fl/fl}$) mice of either sex were used for electrophysiological and imaging experiments. All animals were group housed in a standard 12 h. light/12 h. dark cycle. Animal handling and use followed a protocol approved by the Animal Care and Use Committee of Albert Einstein College of Medicine, in accordance with the National Institutes of Health guidelines. *Bdnf*$^{fl/fl}$ and *TrkB*$^{fl/fl}$ mice were generously donated by Dr. Lisa Monteggia at the University of Texas (Southwestern Medical Center).

#### Presynaptic *Bdnf* and *TrkB* conditional knockout combined to optogenetics

A mix of Cre recombinase-containing adeno-associated viruses AAV5.CamKII.Cre-mCherry (University of Pennsylvania Vector Core) and Cre-dependent ChiEF (AAV-DJ.FLEX.ChiEF-tdTomato) was injected into the DG of 3–4 weeks old (w.o.) *Bdnf*$^{fl/fl}$ or *TrkB*$^{fl/fl}$ mice. WT littermates injected with the same viruses served as controls. The mix of viruses (1 µL total, 1:2 ratio) was injected at a rate of 0.1 µL/min unilaterally into the DG (2.18 mm posterior to bregma, 1.5 mm lateral to bregma, 2.20 mm ventral from dura, for 4.2-mm lambda-bregma distance) of *Bdnf*$^{fl/fl}$, *TrkB*$^{fl/fl}$ and WT mice. All coordinates were adjusted to the lambda-bregma distance for each animal. Briefly, animals were placed on a stereotaxic apparatus (Kopf Instruments) and anesthetized with isoflurane (5% for induction and 1.5% for maintenance). Slices for electrophysiology were prepared from injected animals 4–5 weeks after surgery. This strategy allowed us to optically activate Cre/ChiEF-positive commissural MC axons and record from GCs in the contralateral hippocampus. For each animal, we verified the presence of tdTomato-expressing, putative MC axon terminals in the IML of contralateral slices (fig. S4).

#### Postsynaptic *Bdnf* and *TrkB* conditional knockout

AAV5.CamKII.mCherry (control) or AAV5.CamKII.Cre-mCherry (University of Pennsylvania Vector Core) were injected (1 µL, flow rate of 0.1 µL/min) bilaterally into the upper blade of the DG (2.06 mm posterior to bregma, ± 1.5 mm lateral to bregma, 1.65 mm ventral from dura) of 3–4 w.o. *Bdnf*$^{fl/fl}$ or *TrkB*$^{fl/fl}$ mice. Slices for electrophysiology were prepared from injected animals 10 days after surgery. For each animal, we verified the absence of mCherry-expressing cells in the hilus of the whole ipsilateral hippocampus (fig. S3).

#### Hippocampal slice preparation

Acute transverse hippocampal slices (300-µm thick) were prepared from *Bdnf*$^{fl/fl}$, *TrkB*$^{fl/fl}$ and WT mice, for electrophysiology or imaging experiments. Briefly, the hippocampi were isolated and cut using a VT1200s microslicer (Leica Microsystems Co.) in an ice-cold cutting solution containing (in mM): 110 choline, 2.5 KCl, 25 $NaHCO_3$, 1.25 $NaH_2PO_4$, 0.5 $CaCl_2$, 7 $MgCl_2$, 25 D-glucose, 11.6 sodium L-ascorbate, and 3.1 sodium pyruvate. Slices were then transferred and incubated for 30 min in a chamber placed in a warm bath (33–34°C) with extracellular artificial cerebrospinal fluid (ACSF) solution containing (in mM): 124 NaCl, 2.5 KCl, 26 $NaHCO_3$, 1 $NaH_2PO_4$, 2.5 $CaCl_2$, 1.3 $MgSO_4$ and 10 D-glucose. Slices were kept at room temperature for at least 30 min prior to experiments. All solutions were equilibrated with 95% $O_2$ and 5% $CO_2$ (pH 7.4).

#### Electrophysiology

All recordings were performed at 28 ± 1°C in a submersion-type recording chamber perfused at 2 mL/min with ACSF supplemented with the $GABA_A$ receptor antagonist picrotoxin (100 µM), unless otherwise stated. Whole-cell patch-clamp recordings using a Multiclamp 700A amplifier (Molecular Devices) were made from GCs voltage clamped at −60 mV using patch-type pipette electrodes (~3–4 MΩ) containing (in mM): 131 cesium gluconate, 8 NaCl, 1 CaCl2, 10 EGTA, 10 D-glucose, and 10 HEPES (pH 7.2, 285-290 mOsm). For fig. S5, whole-cell patch-clamp recordings were made using $K^+$-based internal solution, containing the following intracellular solution (in mM): 135 $KMeSO_4$, 5 KCl, 1 $CaCl_2$, 5 NaOH, 10 HEPES, 5 MgATP, 0.4 $Na_3GTP$, 5 EGTA and 10 D-glucose, pH 7.2 (280-290 mOsm). Series resistance (~7–28 MΩ) was monitored throughout all experiments with a −5 mV, 80 ms voltage step, and cells that exhibited a significant change in series resistance (> 20%) were excluded from analysis. All experiments were performed in an interleaved fashion –i.e., control experiments were performed every test experiment on the same day.

A stimulating patch-type pipette filled with ACSF was placed in the IML (< 30 μm from the border of the GC body layer) to activate MC axons. To elicit synaptic responses, paired, monopolar square-wave voltage or current pulses (100–200 μs pulse width, 4–25 V) were delivered through a stimulus isolator (Isoflex, AMPI). Typically, stimulation intensity was adjusted to obtain comparable magnitude synaptic responses across experiments (e.g., 50–100 pA EPSCs at $V_h$ −60 mV). For minimal stimulation (Fig. 5A–C), stimulating pipettes were made from theta glass capillaries. MC-GC LTP was typically induced with brief bursts (5 pulses, 100 Hz) repeated 50 times at 2 Hz (for a total of 25 seconds) while postsynaptic neurons were continuously voltage-clamped at −60 mV. GC firing (fig. S5) was assessed using burst stimulation in the IML (5 pulses at 20 Hz every 20 s), while the membrane potential was held at -65 mV. For this experiment, the recording configuration was switched from voltage-clamp to current-clamp mode.

Electrophysiological data were acquired at 5 kHz, filtered at 2.4 kHz, and analyzed using custom-made software for IgorPro 7.01 (Wavemetrics, Inc.). The magnitude of LTP was determined by comparing 10 min baseline responses with responses 20–30 min after LTP induction or forskolin-induced potentiation. PPR was defined as the ratio of the amplitude of the second EPSC, to the amplitude of the first EPSC. CV was calculated as the standard deviation of EPSC amplitude divided by the mean of EPSC amplitude. Both PPR and CV were measured during 10-min baseline. Averaged traces include 20 consecutive individual responses.

**Optogenetics**

At least 4 weeks post-surgery, acute hippocampal slices were prepared as previously described and slices showing optimal tdTomato expression in the IML of the contralateral DG were used for electrophysiology. Pulses of blue light (0.5−2 ms width duration) were provided using a 473-nm LED (ThorLabs, Inc.) through the microscope objective (40x, 0.8 NA) and centered in the IML.

**Two-photon BDNF-pHluorin imaging**

A mix of Cre recombinase-containing virus (AAV5.CamKII.Cre-mCherry) and Cre-dependent BDNF-pHluorin (AAV-DJ.DIO.BDNF-pHluorin) was injected into the DG of 3–4 weeks old WT or $TrkB^{fl/fl}$ mice. To measure presynaptic BDNF release, the mix of viruses (1 μL total, 1:2 ratio) was injected at a flow rate of 0.1 μL/min unilaterally into the hilus (2.18 mm posterior to bregma, 1.5 mm lateral to bregma, 2.20 mm ventral from dura, for 4.2-mm lambda-bregma distance). To measure postsynaptic BDNF release, the mix was injected bilaterally into the upper GC layer (2.06 mm posterior to bregma, ± 1.5 mm lateral to bregma, 1.65 mm ventral from dura) to avoid expression in MCs (fig. S3). Acute hippocampal slices were prepared as previously described and slices showing optimal mCherry expression were taken for imaging sessions.

All imaging experiments were performed in a submersion-type recording chamber perfused at 2 mL/min with ACSF. Briefly, a broken tip stimulating pipette was placed in the IML at least 100 μm away from the imaging site. Extracellular field recordings were monitored simultaneously using a patch-type pipette filled with 1 M NaCl and placed in IML. The Insight Deep See laser (Spectra Physics) was tuned to 880 nm and the imaging site was selected by the appearance of BDNF-pH puncta in the IML. Using 512 X 512 pixel resolution, the region of interest (ROI) was magnified to 6–8 X and a baseline acquisition of 40 consecutive images at 0.2 Hz was acquired using T-series software (PrairieView 5.4, Bruker Corp.). Following baseline acquisition, a brief MC burst-stimulation (5 pulses at 100 Hz, repeated 50 times at 2 Hz) was delivered and at least 80 consecutive images were acquired at 0.2 Hz. To verify reactivity of the ROI an isosmotic solution of $NH_4Cl$ (50 mM) was added at the end of the imaging session as previously reported (*32, 33*). ROIs were defined manually for each experiment, blind to the injection conditions or treatments. The ROI fluorescence intensity was measured, and the $\Delta F/F_0$ of BDNF-pH signal was determined, using ImageJ software.

**Enriched Environment**

Standard mouse cages (28 cm x 18 cm) containing a feeder and a water dispenser were used as home cages (HC). Enriched environment (EE) housing consists in a large cage (121 cm x 61 cm) containing a feeder, a water dispenser, multiple running wheels, plastic tubes, domes, and other plastic toys (fig. S6A–B). Animals were exposed for two weeks or one hour. In the first case, objects were moved every day to maintain novelty. EE refers to both the effects of cognitive enrichment strictly defined (i.e., exploration, toys, large area), as well as exercise on running wheels. Experimenters were blind to condition when performing and analyzing experiments.

**Electron microscopic immunohistochemistry**

Tissues were collected from male adult (~2–3 months old) Bdnf-HA knock-in mice obtained in a previous study (*34*). Mice were deeply anaesthetized with sodium pentobarbital (150 mg/kg, i.p.) and perfused through the aortic arch with a saline solution containing 2% heparin (~5 mL) followed by a solution (~30 mL) containing 3.75% acrolein and 2% paraformaldehyde in 0.1 M phosphate buffer (pH 7.4; PB). Brains then were sectioned (40-µm thick) on a Vibratome, collected in PB and transferred in a cryoprotectant solution containing 30% sucrose and 30% ethylene glycol in PB and stored at -20°C. As previously described (*34*), sections were processed for HA-immunoperoxidase labelling and flat embedding in EMbed-812 (EMS) between two sheets of Aclar plastic. Sections containing the DG were selected from the plastic embedded sections and glued onto Epon chucks and trimmed to 1-mm trapezoids. Ultra-thin sections (70-nm thick) through the tissue-plastic interface were cut with a diamond knife (EMS) on a Leica EM UC6 ultratome, and sections were collected on 400-mesh, thin-bar copper grids (EMS). Grids were then counterstained with uranyl acetate and Reynold's lead citrate. Ultra-thin sections were then examined on a Tecnai Biotwin transmission electron microscope (FEI) equipped with an AMT digital camera. Cell profiles were identified by defined morphological criteria (*58*). Dendritic profiles generally were postsynaptic to axon terminals and contained regular microtubule arrays, whereas dendritic spines also were usually postsynaptic to axon terminal profiles. Axon terminals contained small synaptic vesicles and dense-core vesicles. HA-immunoperoxidase labelling was evident as a characteristic, electron-dense DAB reaction product precipitate.

**Reagents**

Reagents were bath applied following dilution into ACSF from stock solutions stored at −4°C or −20°C prepared in water or DMSO, depending on the manufacturer's recommendation. The final DMSO concentration was < 0.1% total volume. Reagents *N*-(piperidin-1-yl)-5-(4-iodophenyl)-1-(2,4-dichlorophenyl)-4-methyl-1*H*-pyrazole-3-carboxamide (AM251), (*R*)-(+)-[2,3-Dihydro-5-methyl-3-(4-morpholinylmethyl)pyrrolo[1,2,3-*de*]-1,4-benzoxazin-6-yl]-1-naphthalenylmethanone mesylate (WIN 55,212-2), *N*-[2-[[(Hexahydro-2-oxo-1*H*-azepin-3-yl)amino]carbonyl]phenyl]benzo[b]thiophene-2-carboxamide (ANA-12), forskolin, cyclopiazonic acid, U73122 (1-[6-[[(17β)-3-Methoxyestra-1,3,5(10)-trien-17-yl]amino]hexyl]-1H-pyrrole-2,5-

dione) and picrotoxin were purchased from Tocris Bioscience. Dimethyl Sulfoxide (DMSO) and cadmium were purchased from Sigma-Aldrich. Recombinant human BDNF protein was purchased from Hello Bio. Salts for making ACSF and internal solutions were purchased from Sigma-Aldrich.

**Quantification and Statistical Analysis**

Statistical analysis was performed using OriginPro software (OriginLab version b9.2.272). For imaging, analysis was done blind to the conditions. The normality of distributions was assessed using the Shapiro-Wilk test. In normal distributions, Student's unpaired and paired two-tailed t-tests were used to assess between-group and within-group differences, respectively. The non-parametric paired sample Wilcoxon signed rank test and Mann-Whitney's U test were used in non-normal distributions. Statistical significance was set to $p < 0.05$ (*** indicates $p < 0.001$, ** indicates $p < 0.01$, and * indicates $p < 0.05$).

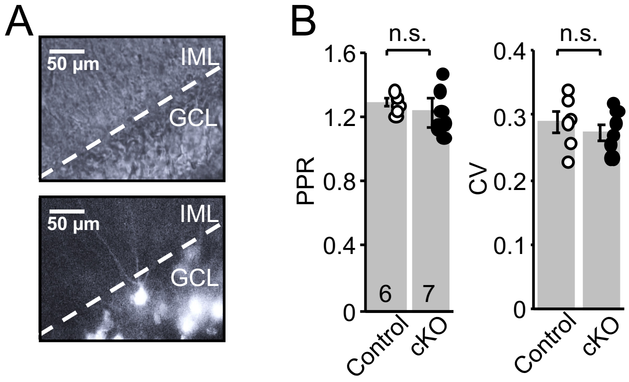


**Fig. S1 (related to Fig. 3). Postsynaptic *Bdnf* deletion does not significantly affect MC-GC presynaptic properties.**
**(A)** Infrared/differential interference contrast (*top*) and fluorescence (*bottom*) images showing mCherry was sparsely expressed in the upper GC layer of $Bdnf^{fl/fl}$. **(B)** Summary plots showing PPR (Control: 1.3 ± 0.02, n = 6; Post *Bdnf* cKO: 1.2 ± 0.05, n = 7; Control vs cKO, p = 0.3813, unpaired t-test) and CV (Control: 0.29 ± 0.02, n = 6; Post *Bdnf* cKO: 0.27 ± 0.01, n = 7; Control vs cKO, p = 0.4522, unpaired t-test) were unchanged in Post *Bdnf* cKO mice as compared to controls.

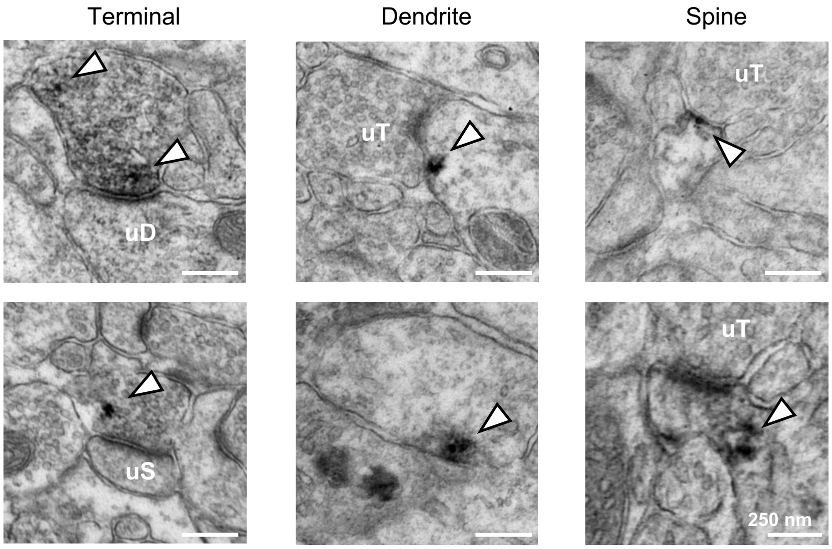


**Fig. S2 (related to Fig. 3). BDNF localizes to MC axon terminals, dendrites, and dendritic spines of GCs.**
White arrows indicate HA-immunoperoxidase labelling in the IML from Bdnf-HA mice (N = 2), visualized by electron microscopy. uD: unlabeled dendrite; uT: unlabeled terminal; uS: unlabeled spine.

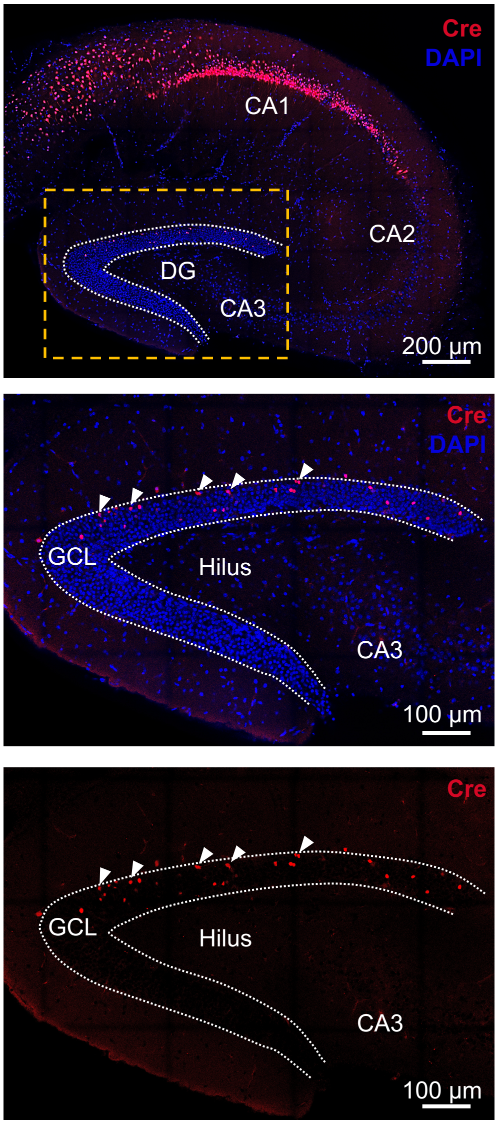


**Figure S3 (related to Fig. 3). Cre is restricted to the upper GC layer.**
Confocal images of Cre-mCherry expression in the DG of AAV5.CamKII.Cre-mCherry-injected control mice (*Top*). High magnification (*Middle and Bottom*) showing that GCs are selectively infected with Cre-mCherry (white arrowheads).

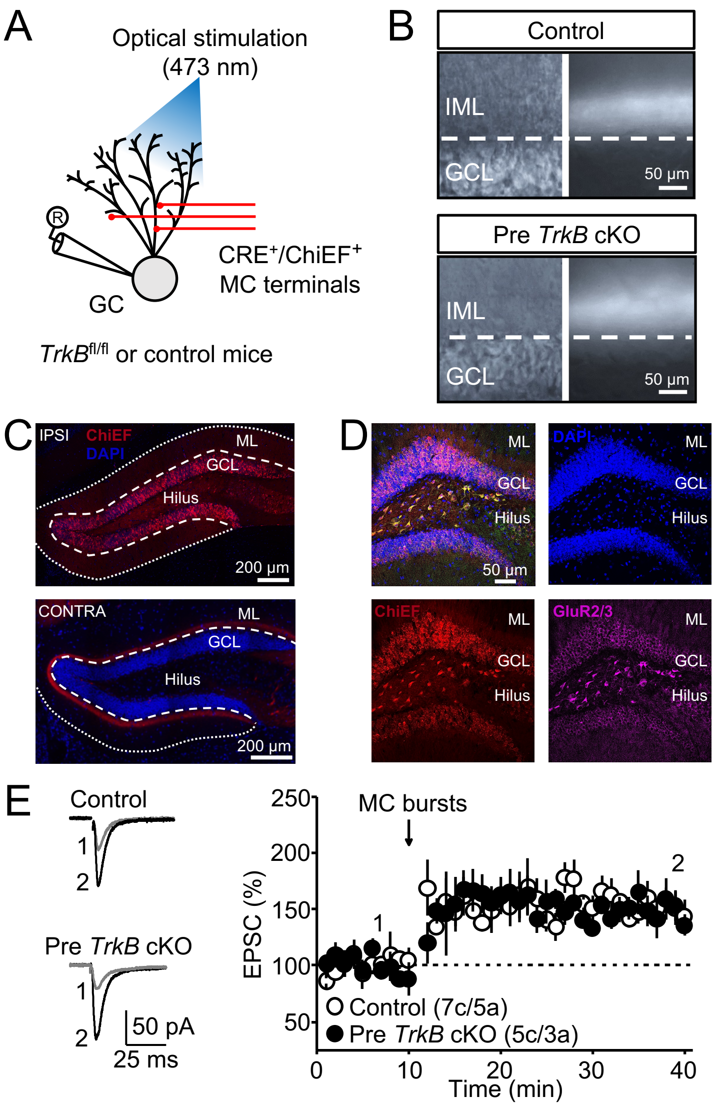


**Fig. S4 (related to Fig. 3). Presynaptic TrkB is not required for MC-GC LTP.**
**(A)** Schematic diagram illustrating the optical stimulation of ChIEF-expressing MC axons and the recording configuration of GCs in contralateral hippocampal slices. A mix of AAV5.CaMKII.Cre-mCherry and AAV-DJ.FLEX.ChIEF-tdTomato was injected in the hilus of 3–4 w.o. WT (control) or *TrkB*$^{fl/fl}$ (Pre *TrkB* cKO) littermates. Whole-cell patch-clamp recordings were performed from contralateral GCs. **(B)** Infrared/differential interference contrast (*left*) and fluorescence (*right*) images showing ChIEF-tdTomato was selectively expressed in the IML of contralateral DG of control and Pre *TrkB* cKO mice. **(C–D)** Confocal images showing the expression of ChiEF-tdTomato in the injection site (ipsi) and in the IML of the contralateral DG of *TrkB*$^{fl/fl}$ mice. **(D)** Confocal images of ChIEF-tdTomato expression in hilar MCs labelled with the GluR2/3 marker in the ipsilateral DG. **(E)** Optically-induced LTP was intact in Pre *TrkB* cKO mice as compared to controls (Control: 152 ± 11% of baseline, n = 7, p < 0.01, paired t-test; Pre *TrkB* cKO: 159 ± 12% of baseline, n = 5, p < 0.01, paired t-test; Control vs cKO, p = 0.6767, unpaired t-test). Arrows correspond to the burst-stimulation of MC axons. Representative traces (1) and (2) correspond to the areas (1) and (2) in the summary time-course plots. Data are represented as mean ± SEM. Number of cells and mice are showed between parentheses.

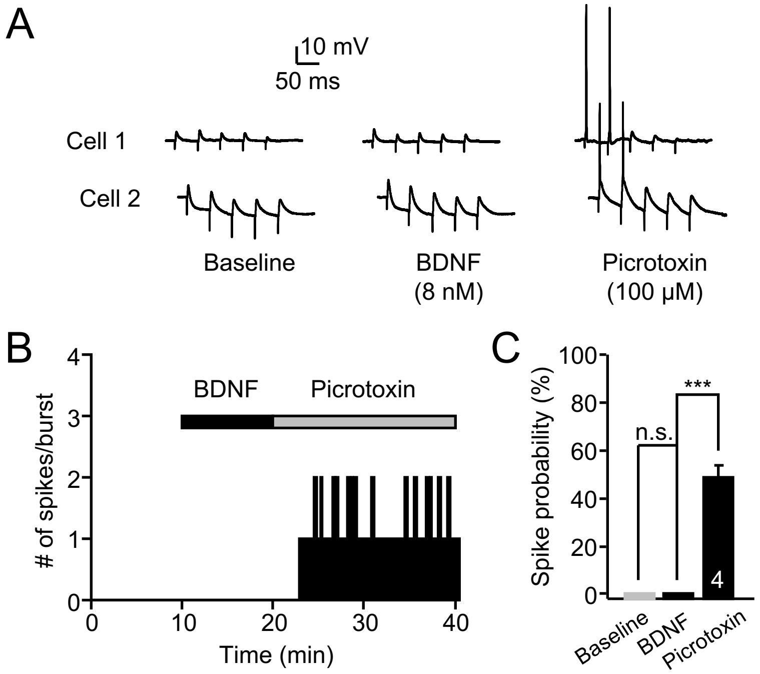


**Fig. S5 (related to Fig. 4). Bath application of BDNF alone does not trigger GC firing.**
**(A)** Sample traces of two representative experiments showing burst stimulation before and after application of BDNF (8 nM) or picrotoxin (100 μM). **(B)** Time-course plot (representative experiment) of the number of spikes per burst following bath application of 8 nM BDNF and 100 μM picrotoxin. **(C)** Summary data showing the spike probability after bath application of BDNF and picrotoxin in the same GC (spike probability in picrotoxin: 49 ± 5%, n = 4; baseline vs picrotoxin, $p < 0.01$, one-way ANOVA followed by a Tukey's *post hoc* test).

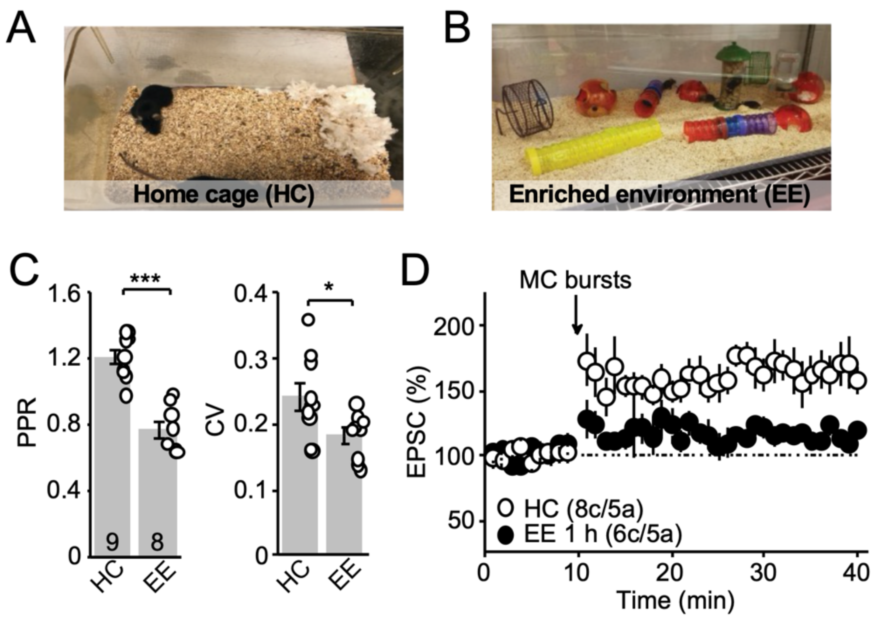


**Fig. S6 (related to Fig. 5). Short exposure to enriched environment induces MC-GC strengthening.**
**(A–B)** Images showing the enriched environment cage (EE) and home cage (HC). **(C)** Summary plots showing that PPR and CV were decreased in mice following a 1-hour exposure (PPR HC: 1.2 ± 0.05, n = 9; EE: 0.8 ± 0.05, n = 8; HC vs EE, $p < 0.001$, unpaired t-test; CV HC: 0.24 ± 0.02, n = 9; EE: 0.18 ± 0.01, n = 8; HC vs EE, $p < 0.05$, unpaired t-test). **(D)** Synaptically-induced LTP was decreased in mice following 1-hour exposure to EE as compared to controls (HC: 165 ± 14% of baseline, n = 8, $p < 0.01$, Wilcoxon signed rank test; EE: 115 ± 2% of baseline, n = 6, $p < 0.001$, paired t-test; HC vs EE: $p < 0.01$, Mann-Whitney's U test). Data are represented as mean ± SEM. *** $p < 0.001$; * $p < 0.05$.

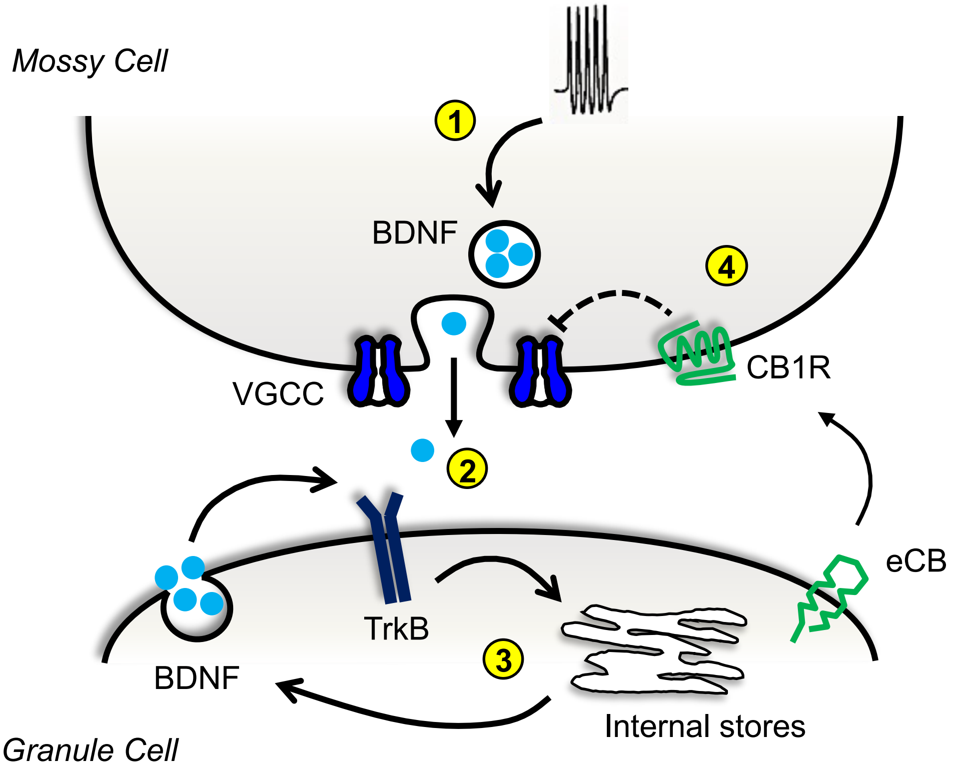


**Fig. S7 (related to Fig. 1–5). Working model.**
Repetitive activity of MC axons releases BDNF, which by activating postsynaptic TrkB likely activates PLC and mobilizes calcium internal stores, thereby inducing postsynaptic BDNF secretion (i.e., from GCs) and long-lasting MC-GC synaptic strengthening. In addition, $CB_1R$ activity tightly inhibits presynaptic BDNF release, thereby controls MC-GC synaptic strengthening.
